## Supplementary material for "A small molecule inhibitor of PTP1B and PTPN2 enhances T cell anti-tumor immunity": SUPP FIGS 1-13

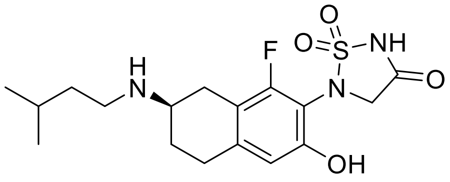


ABBV-CLS-484

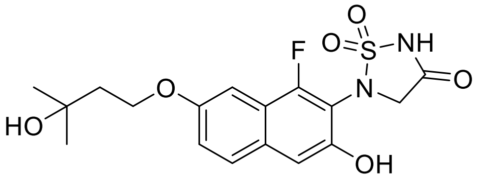


Compound 182

***Figure S1. The chemical structures of ABBV-CLS-484 and Compound 182.***

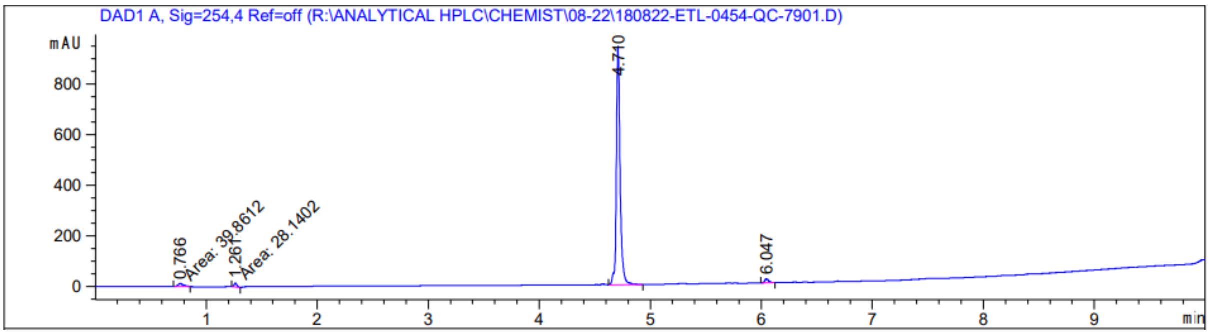


| Peak # | RetTime [min] | Type | Width [min] | Area [mAU*s] | Height [mAU] | Area % |
|---|---|---|---|---|---|---|
| 1 | 0.766 | MM | 0.0549 | 39.86118 | 12.11183 | 1.7379 |
| 2 | 1.261 | MM | 0.0280 | 28.14021 | 16.72783 | 1.2269 |
| 3 | 4.710 | BV | 0.0351 | 2189.17505 | 942.49902 | 95.4455 |
| 4 | 6.047 | BB | 0.0364 | 36.46122 | 15.54548 | 1.5897 |

***Figure S2. HPLC Chromatogram of Compound 182 and Purity.*** HPLC chromatogram of XI (Compound 182; absorbance measured at wavelength 254 nm) and purity measurement. The compound had a retention time of 4.710 min and a purity of 95% based on peak area (area under the curve).

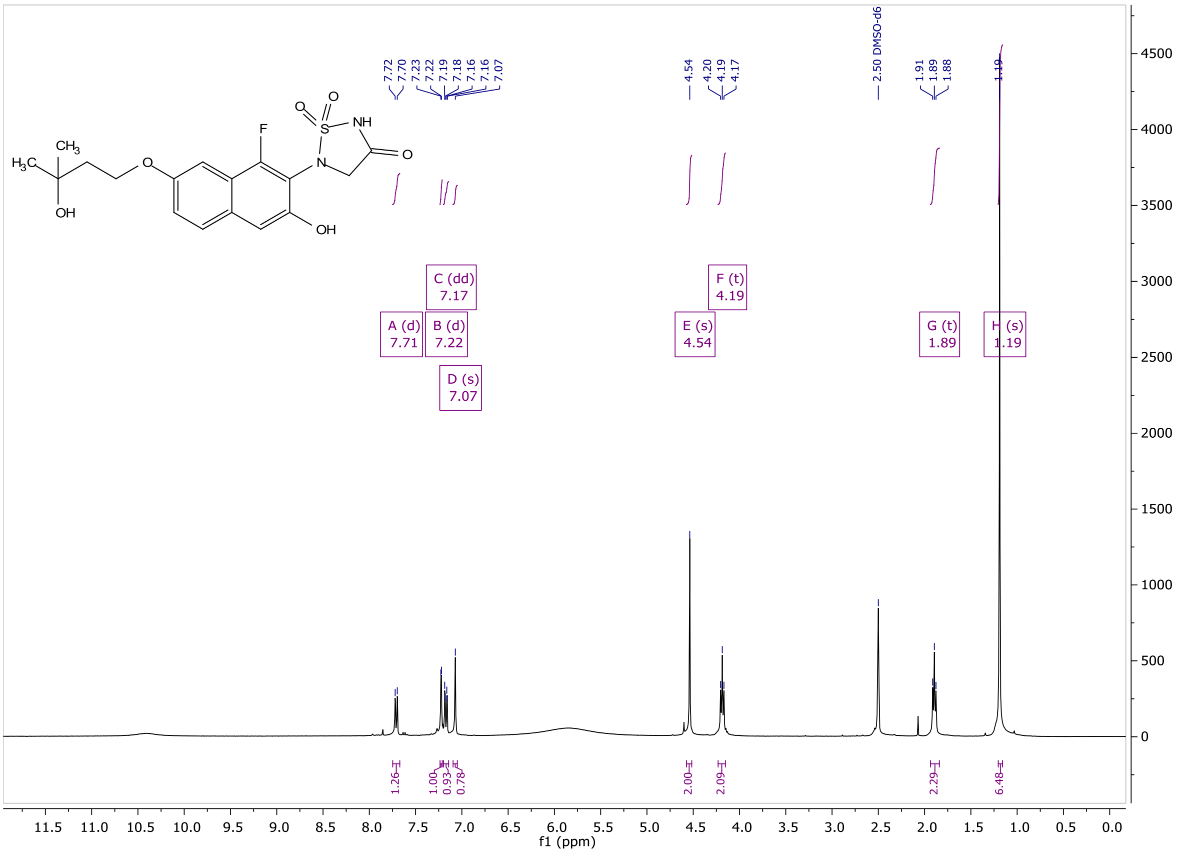


***Figure S3.*** ***[1]H NMR spectrum of Compound 182.*** [1]H NMR spectrum of XI (Compound 182) obtained in deuterated DMSO-$d_6$ at 400 MHz.

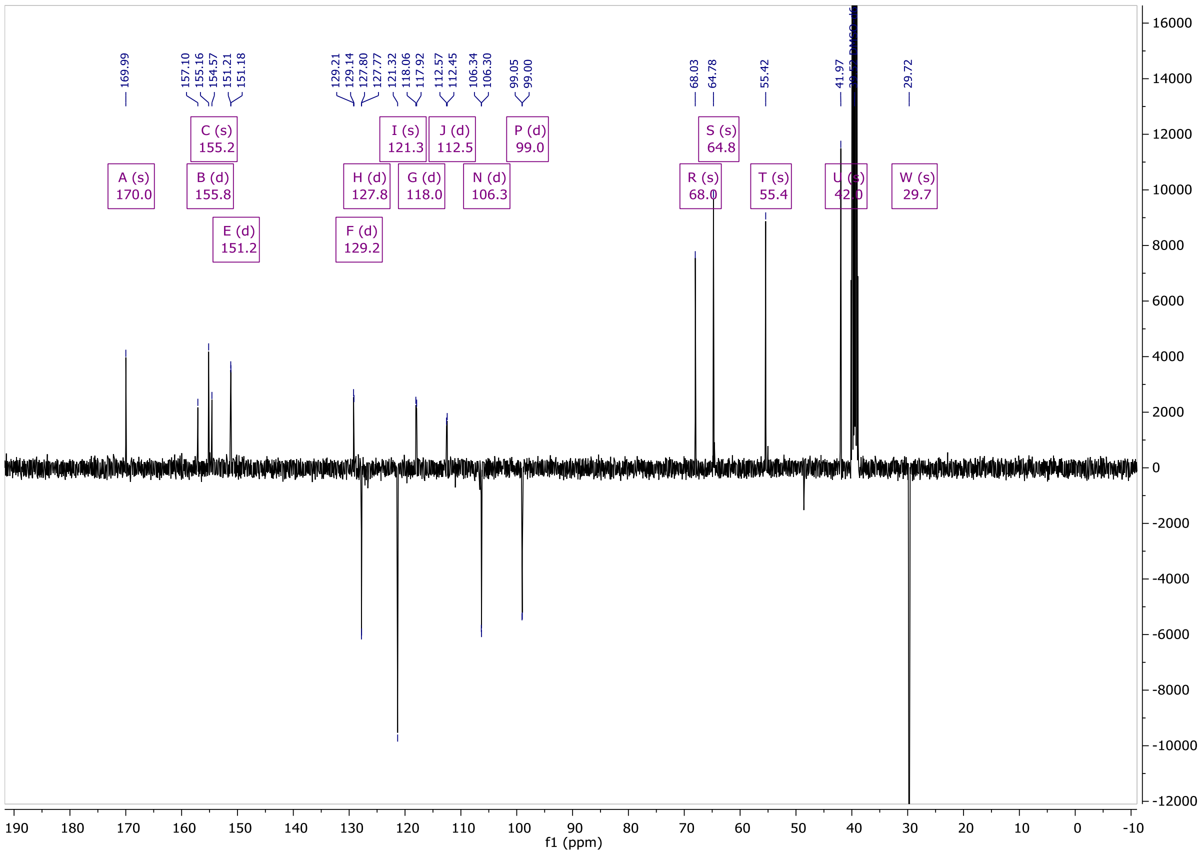


***Figure S4.*** ***[13]C NMR spectrum of Compound 182.*** $^{13}$C-DEPTQ NMR spectrum of XI (Compound 182) obtained in deuterated DMSO-$d_6$ at 101 MHz.

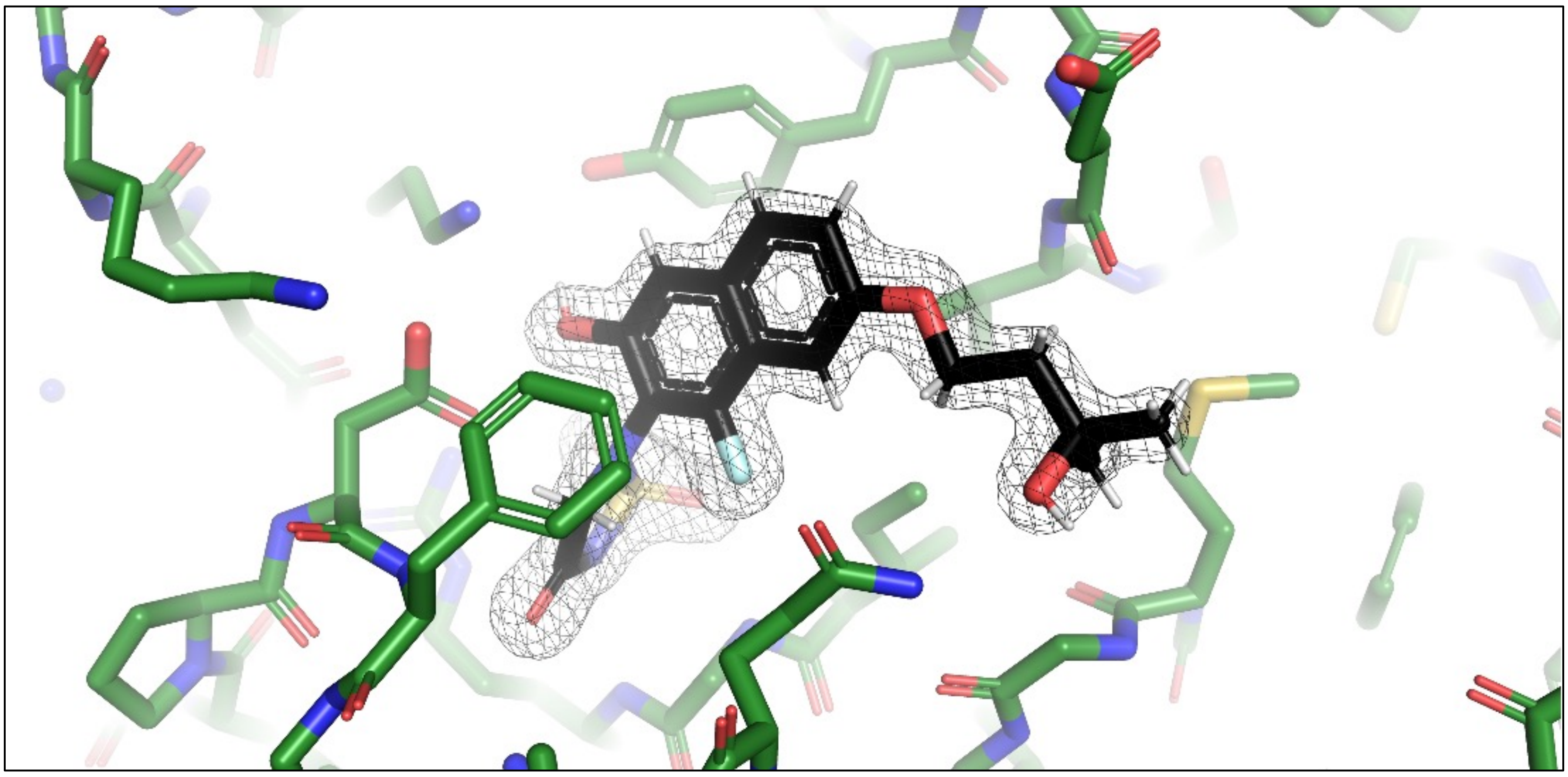

**Fig. S5**

***Figure S5. Electron density for Compound 182.*** The region about the compound in the PTP1B/Compound 182 complex 2Fo-Fc electron density map is shown as a mesh and is contoured at 2σ. See Table 2 for crystallographic details.

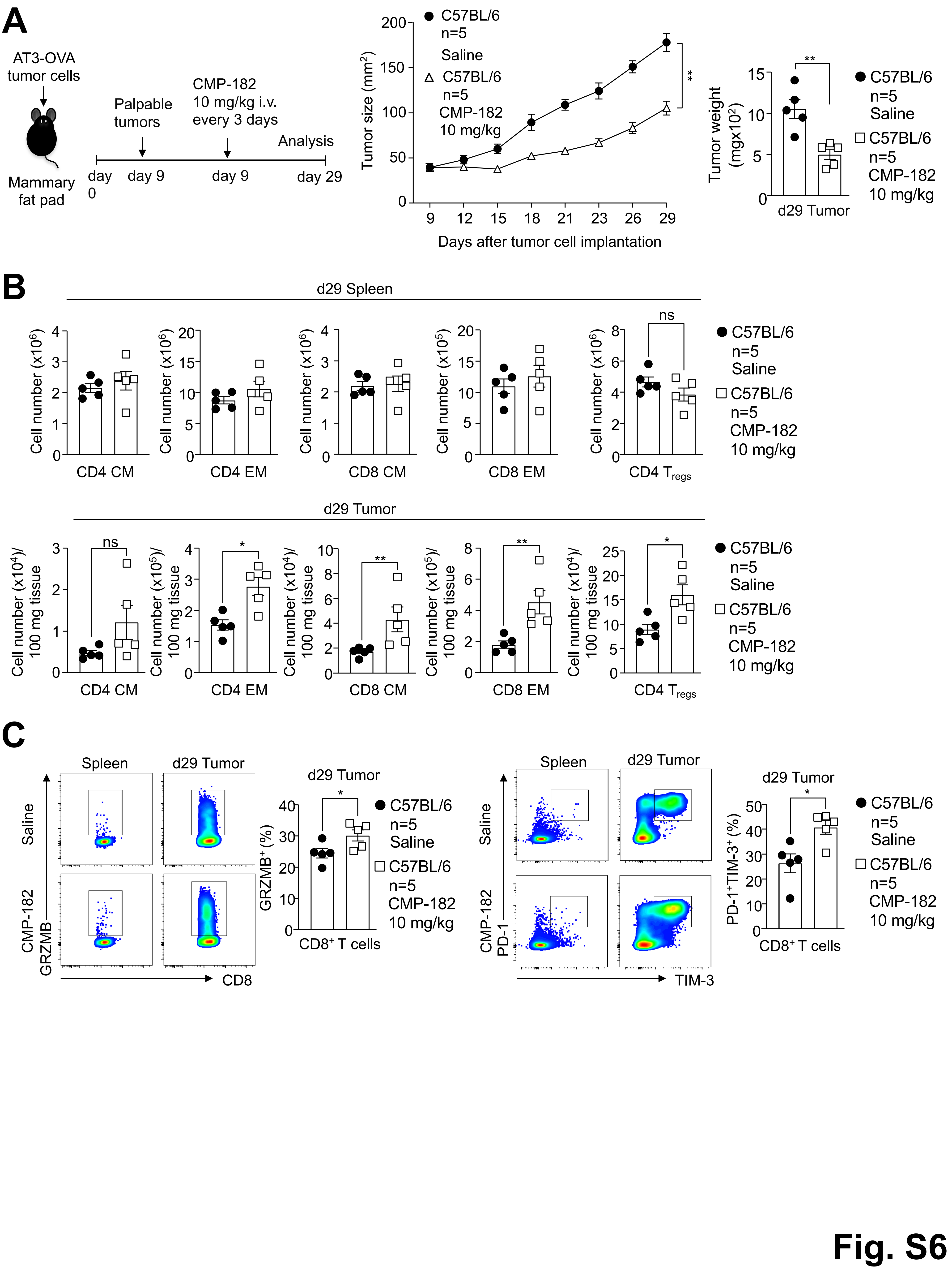
A
AT3-OVA tumor cells
Mammary fat pad
Palpable tumors
CMP-182 10 mg/kg i.v. every 3 days
Analysis
day 0
day 9
day 9
day 29
Tumor size (mm²)
Days after tumor cell implantation
C57BL/6 n=5 Saline
C57BL/6 n=5 CMP-182 10 mg/kg
Tumor weight (mgx10²)
d29 Tumor
B
d29 Spleen
Cell number (x10⁶)
Cell number (x10⁵)
CD4 CM
CD4 EM
CD8 CM
CD8 EM
CD4 Tregs
ns
d29 Tumor
Cell number (x10⁴)/ 100 mg tissue
Cell number (x10⁵)/ 100 mg tissue
C
Spleen
d29 Tumor
Saline
CMP-182
GRZMB
CD8
GRZMB⁺ (%)
CD8⁺ T cells
PD-1
TIM-3
PD-1⁺TIM-3⁺ (%)
Fig. S6

***Figure S6. Administration of Compound 182 every three days represses tumor growth.*** AT3-OVA mammary tumor cells were injected into the fourth inguinal mammary fat pads of C57BL/6 mice. Mice were treated with Compound 182 (10 mg/kg i.v.) or saline on days 9, 12, 15, 18, 21, 24 and 27 after tumor cell implantation. **a**) Tumor growth was monitored and tumor weights measured. **b**) Tumor-infiltrating lymphocytes (TILs) or splenocytes including $CD44^{hi}CD62L^{hi}$ $CD8^{+}$ and $CD4^{+}$ central/memory (CM) T cells and $CD44^{hi}CD62L^{lo}$ $CD8^{+}$ and $CD4^{+}$ effector/memory (EM) T cells and $CD4^{+}CD25^{+}FoxP3^{+}$ regulatory T cells ($T_{regs}$) were analysed by flow cytometry. **c**) Tumor-infiltrating T cells from (a) were assessed for intracellular granzyme B (GrzB) and surface PD-1 and TIM-3 in unstimulated tumor-infiltrating $CD8^{+}$ T cells. Representative results (means ± SEM) from at least two independent experiments are shown. Significance for tumor sizes in (a) was determined using a 2-way ANOVA Test and for tumor weights in (a) using a 2-tailed Mann-Whitney U Test. In (b-c) significances were determined using a 2-tailed Mann-Whitney U Test.

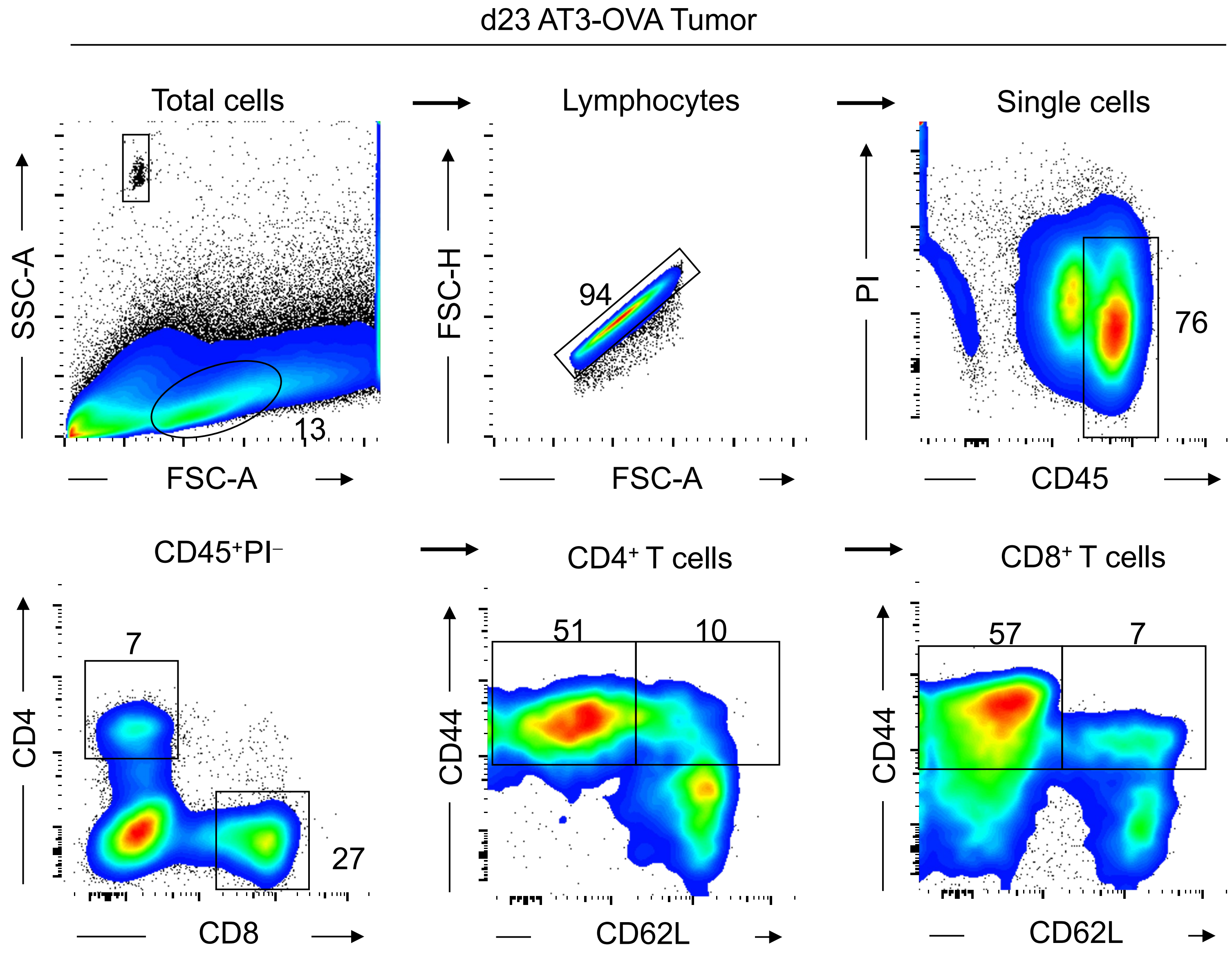

d23 AT3-OVA Tumor
Total cells
Lymphocytes
Single cells
SSC-A
FSC-A
13
FSC-H
FSC-A
94
PI
CD45
76
CD45⁺PI⁻
CD4⁺ T cells
CD8⁺ T cells
CD4
CD8
7
27
CD44
CD62L
51
10
CD44
CD62L
57
7


**Fig. S7**

***Figure S7. Gating strategy for intratumoral lymphocytes.*** AT3-OVA mammary tumor cells were injected into the fourth inguinal mammary fat pads of C57BL/6 mice. Mice were treated with Compound 182 (10 mg/kg i.v.) or saline on days 6, 8, 10, 12, 14, 16, 18 and 21 after tumor cell implantation. Tumor-infiltrating lymphocytes (TILs) including $CD44^{hi}CD62L^{lo}$ $CD8^{+}$ and $CD4^{+}$ effector/memory (EM) T cells were analysed by flow cytometry.

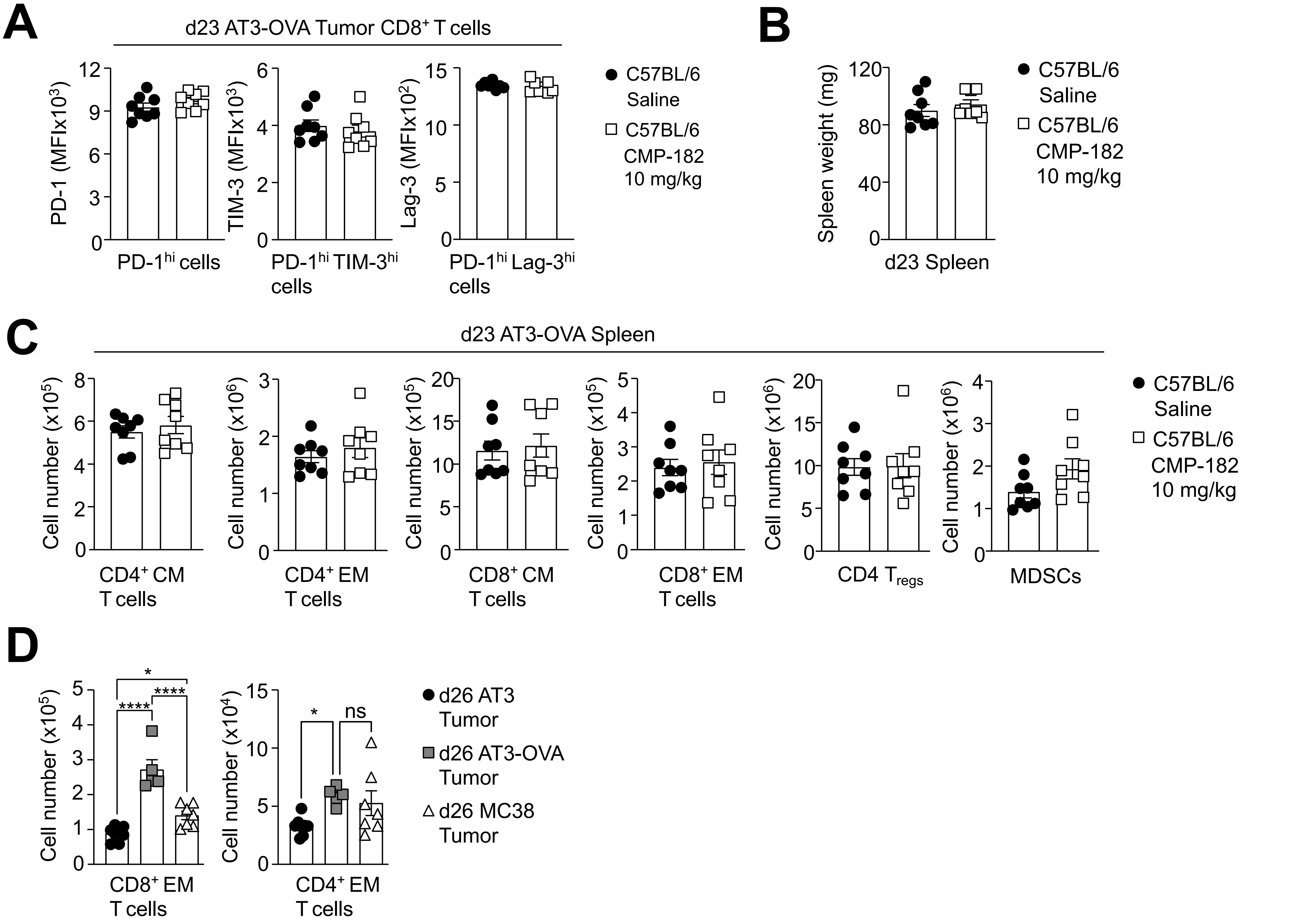
A
d23 AT3-OVA Tumor CD8+ T cells
PD-1 (MFIx10^3)
PD-1^hi cells
TIM-3 (MFIx10^3)
PD-1^hi TIM-3^hi cells
Lag-3 (MFIx10^2)
PD-1^hi Lag-3^hi cells
C57BL/6 Saline
C57BL/6 CMP-182 10 mg/kg
B
Spleen weight (mg)
d23 Spleen
C
d23 AT3-OVA Spleen
Cell number (x10^5)
CD4+ CM T cells
Cell number (x10^6)
CD4+ EM T cells
CD8+ CM T cells
CD8+ EM T cells
CD4 Tregs
MDSCs
D
Cell number (x10^5)
CD8+ EM T cells
Cell number (x10^4)
CD4+ EM T cells
d26 AT3 Tumor
d26 AT3-OVA Tumor
d26 MC38 Tumor
ns

Fig. S8

***Figure S8. Compound 182 does not affect splenic T cell numbers.*** **a-c**) AT3-OVA mammary tumor cells were injected into the fourth inguinal mammary fat pads of C57BL/6 mice. Mice were treated with Compound 182 (10 mg/kg i.v.) or saline on days 6, 8, 10, 12, 14, 16, 18 and 21 after tumor cell implantation. **a**) PD-1, TIM-3 and Lag-3 mean fluorescence intensities (MFI) on PD-1, TIM-3 and Lag-3 on PD-$1^{hi}$ or PD-$1^{hi}$TIM-$3^{hi}$ or PD-$1^{hi}$Lag-$3^{hi}$ tumor-infiltrating (TILs) $CD8^+$ T cells were analysed by flow cytometry. **b**) Spleen weights from mice in (a) were determined. **c**) Splenocytes from mice in (a) including $CD44^{hi}CD62L^{hi}$ $CD8^+$ and $CD4^+$ central/memory (CM) T cells and $CD44^{hi}CD62L^{lo}$ $CD8^+$ and $CD4^+$ effector/memory (EM) T cells, $CD4^+CD25^+FoxP3^+$ regulatory T cells ($T_{regs}$) and granulocytic and monocytic $CD11b^+F4/80^{hi/lo}Ly6C^+Ly6G^{+/-}$ (MDSCs) myeloid-derived suppressor cells were analysed by flow cytometry. **d**) AT3 or AT3-OVA mammary tumor cells were injected into the fourth inguinal mammary fat pads of C57BL/6 mice or MC38 were xenografted (s.c) into the flanks of C57BL/6 mice. $CD8^+$ and $CD4^+$ effector/memory (EM) T cells were analysed by flow cytometry. Representative results (means ± SEM) from at least two independent experiments are shown. In (d) significances were determined using a 1-way ANOVA Test.

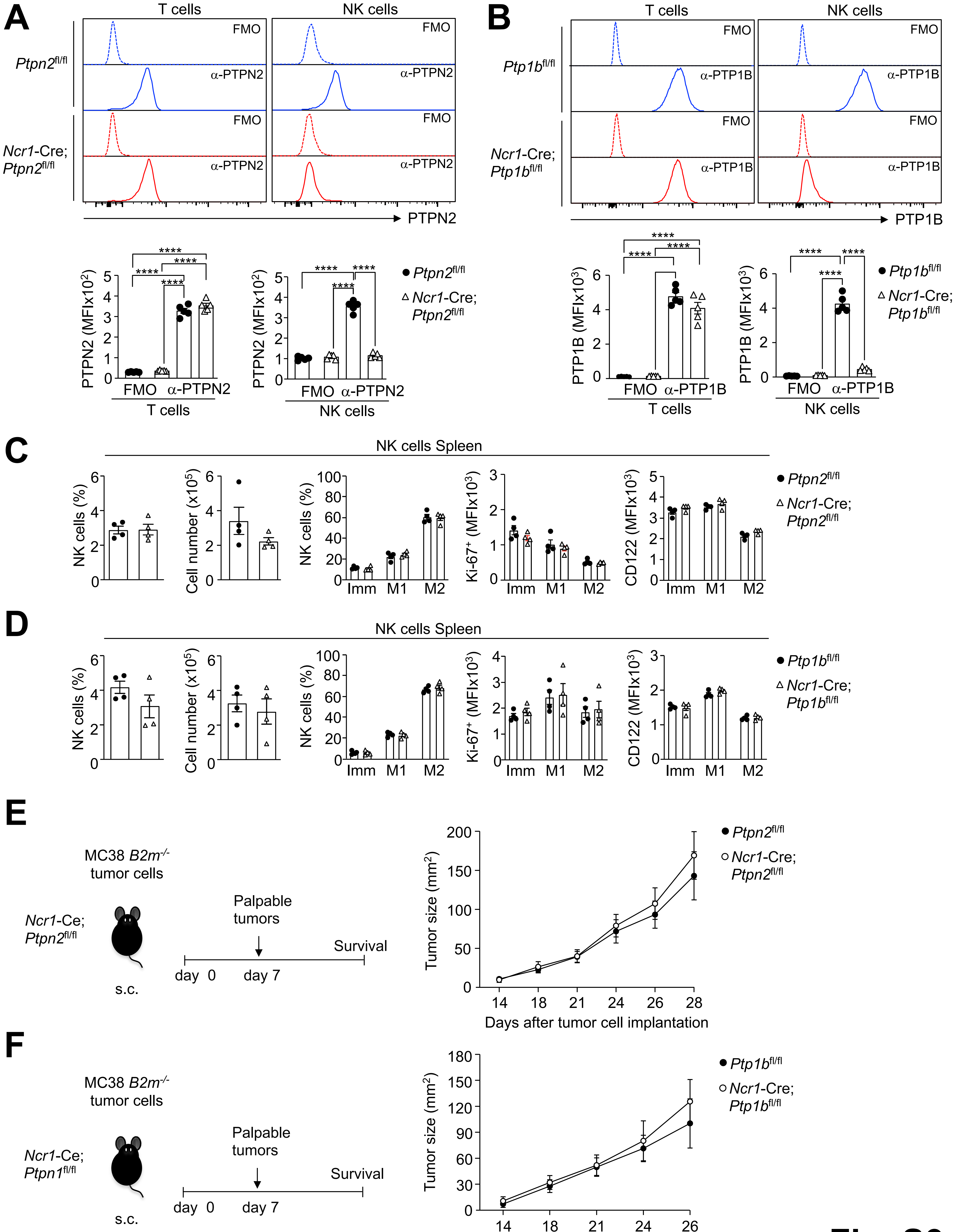
A
T cells
NK cells
Ptpn2fl/fl
Ncr1-Cre; Ptpn2fl/fl
FMO
α-PTPN2
PTPN2
PTPN2 (MFIx10²)
****
FMO α-PTPN2
T cells
NK cells
● Ptpn2fl/fl
△ Ncr1-Cre; Ptpn2fl/fl
B
Ptp1bfl/fl
Ncr1-Cre; Ptp1bfl/fl
α-PTP1B
PTP1B
PTP1B (MFIx10³)
FMO α-PTP1B
● Ptp1bfl/fl
△ Ncr1-Cre; Ptp1bfl/fl
C
NK cells Spleen
NK cells (%)
Cell number (x10⁵)
Ki-67⁺ (MFIx10³)
CD122 (MFIx10³)
Imm M1 M2
D
NK cells Spleen
E
MC38 B2m-/- tumor cells
Ncr1-Ce; Ptpn2fl/fl
s.c.
Palpable tumors
Survival
day 0
day 7
Tumor size (mm²)
Days after tumor cell implantation
● Ptpn2fl/fl
○ Ncr1-Cre; Ptpn2fl/fl
F
MC38 B2m-/- tumor cells
Ncr1-Ce; Ptpn1fl/fl
● Ptp1bfl/fl
○ Ncr1-Cre; Ptp1bfl/fl

**Fig. S9**

***Figure S9. NK cell-specific deletion of PTP1B or PTPN2 does not alter NK cell development and tumor growth.*** **a-b**) Splenic $CD3^+CD19^-CD49b^-NK.1.1^-$ T cells and $CD3^-CD19^-CD49b^{hi}NK.1.1^{hi}$ NK cells from **a**) *Ncr*-Cre;*Ptpn2*$^{fl/fl}$ or **b**) *Ncr*-Cre;*Ptpn1b*$^{fl/fl}$ mice were assessed for PTPN2 and PTP1B protein levels by flow cytometry; FMO staining is background staining. **c-d**) Proportions of NK cells ($CD45^+NK1.1^+$, $CD49b^+NKp46^+CD3^-TCR\beta^-CD19^-F4/80^-$) from **c**) *Ncr*-Cre;*Ptpn2*$^{fl/fl}$ or **d**) *Ncr*-Cre;*Ptpn1b*$^{fl/fl}$ mice in each maturation subset [immature (Imm. $CD27^+$, $CD11b^-$), M1 ($CD27^+$, $CD11b^+$), M2 ($CD27^-$, $CD11b^+$)] and those staining for Ki67 (a marker of proliferation) or CD122 (IL2Rβ/IL15Rβ) were determined by flow cytometry. **e-f**) MC38 *B2m*$^{-/-}$ were xenografted (s.c) into the flanks of **e**) *Ncr*-Cre;*Ptpn2*$^{fl/fl}$ or **f**) *Ncr*-Cre;*Ptpn1b*$^{fl/fl}$ mice and tumor growth was monitored. Representative results from at least two independent experiments are shown. In (a) significances were determined using a 1-way ANOVA Test.

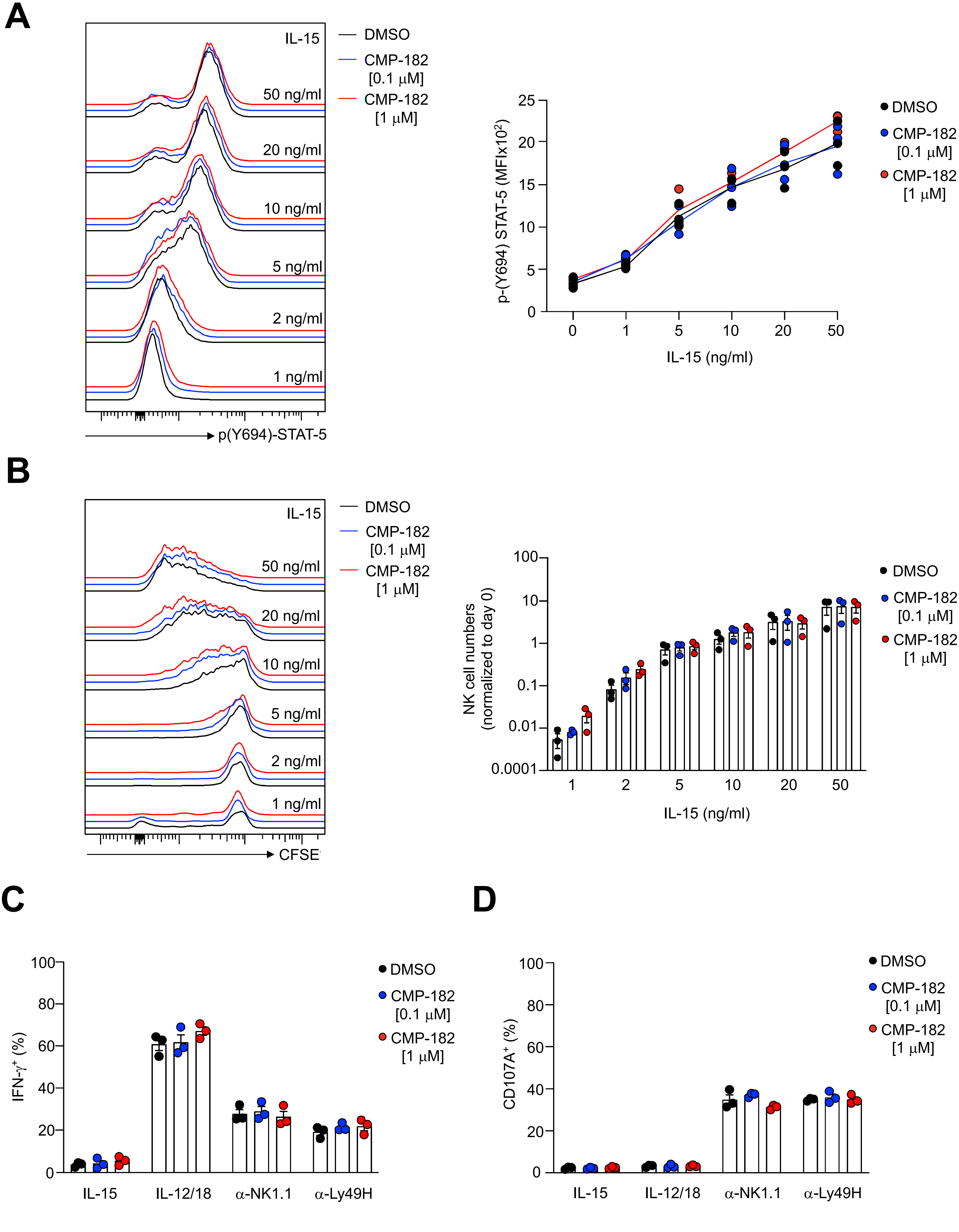
A
IL-15
DMSO
CMP-182 [0.1 μM]
CMP-182 [1 μM]
50 ng/ml
20 ng/ml
10 ng/ml
5 ng/ml
2 ng/ml
1 ng/ml
p(Y694)-STAT-5
p-(Y694) STAT-5 (MFIx10²)
IL-15 (ng/ml)
B
CFSE
NK cell numbers (normalized to day 0)
C
IFN-γ⁺ (%)
IL-15
IL-12/18
α-NK1.1
α-Ly49H
D
CD107A⁺ (%)

Fig. S10

***Figure S10. Compound 182 does not enhance NK cell cytotoxicity and proliferation in vitro.*** **a-b**) NK cells were expanded *ex vivo* in the presence of IL-15 and then pre-treated with Compound 182 (0.1 or 1 µg/ml) or vehicle (1% v/v DMSO) in the absence of IL-15. **a**) NK cells were re-stimulated for 15 min with the indicated amounts of IL-15 and the MFI of Y694 phosphorylated STAT-5 (pSTAT-5) mean fluorescence intensities (MFIs) determined by flow cytometry. **b**) Alternatively, NK cells were stained with carboxyfluorescein succinimidyl ester (CFSE, Invitrogen) and then re-stimulated with the indicated amounts of IL-15 for 96 h and NK cell proliferation (CFSE dilution) monitored by flow cytometry. **c-d**) NK cells were expanded *ex vivo* in the presence of IL-15 and then pre-treated with Compound 182 (0.1 or 1 µg/ml) or vehicle in the absence of IL-15. NK cells were re-stimulated with either IL-12 (10 ng/ml) or IL-18 (100 ng/ml) or with α-NK1.1 or α-Ly49H for 4 h. NK cells were maintained in 10 ng/ml IL-15 throughout the 4 h incubation time. Percentages of **c**) IFN-$\gamma^+$ and **d**) CD107A$^+$ NK cells were determined by flow cytometry. Representative results in (b-d) (means ± SEM) and in (a) presented as line graphs with individual data points from at least two independent experiments are shown.

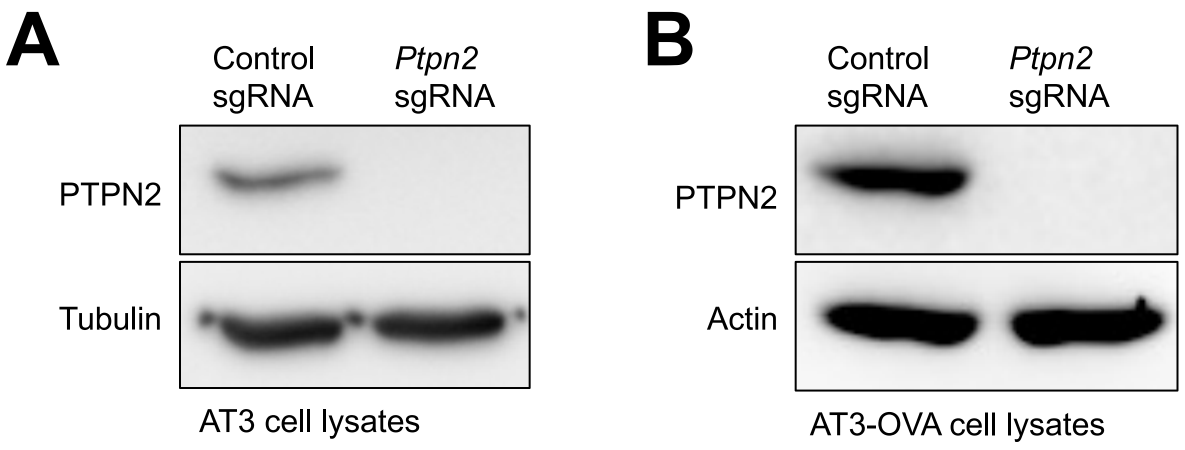
A
Control sgRNA
*Ptpn2* sgRNA
PTPN2
Tubulin
AT3 cell lysates
B
Control sgRNA
*Ptpn2* sgRNA
PTPN2
Actin
AT3-OVA cell lysates

**Fig. S11**

***Figure S11. CRISPR/Cas9-mediated PTPN2 deletion in AT3 and AT3-OVA tumor cells.*** PTPN2 was deleted in **a**) AT3 or **b**) AT3-OVA mammary tumor cells using control or *Ptpn2* sgRNAs and CRISPR RNP gene-editing. Cells were lysed and proteins resolved by SDS-PAGE and immunoblotted for PTPN2 (6F3) and either tubulin or actin.

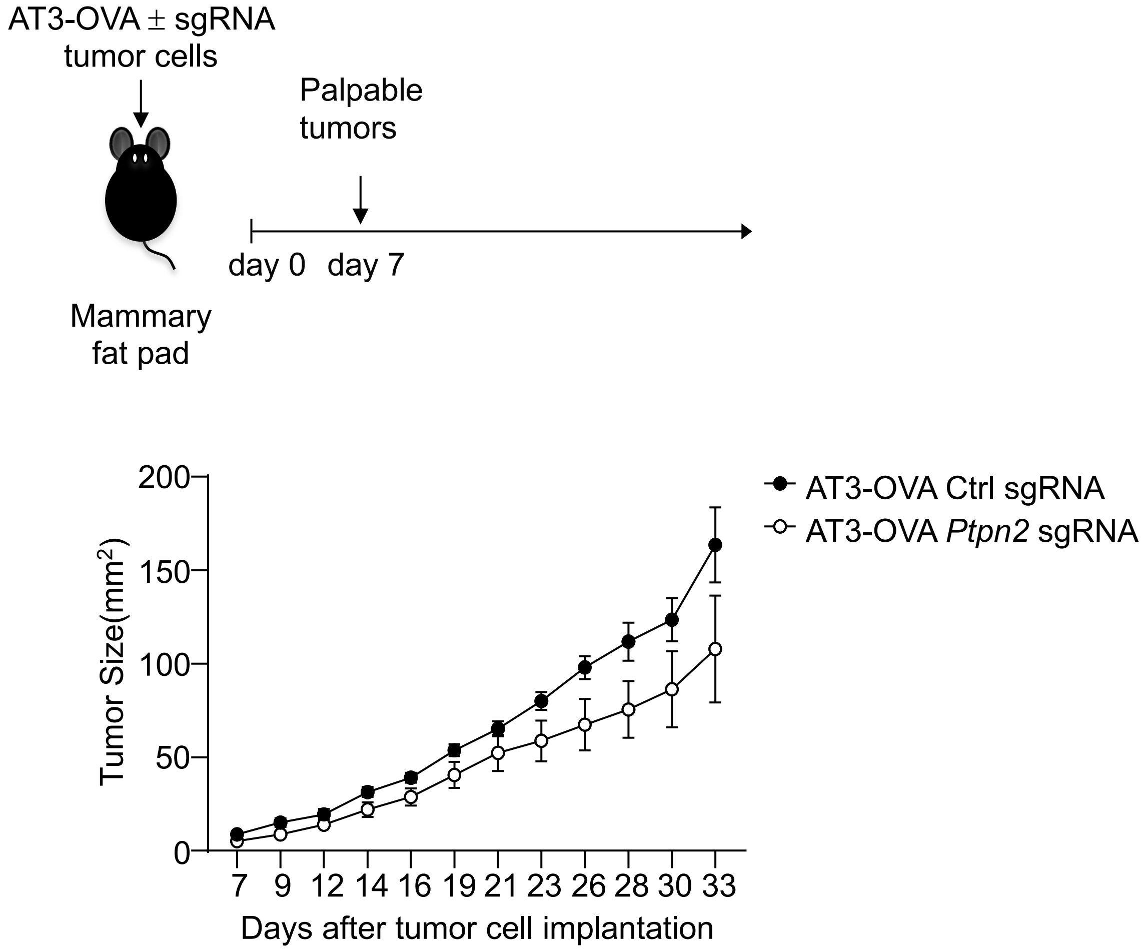
AT3-OVA ± sgRNA
tumor cells
Palpable
tumors
day 0
day 7
Mammary
fat pad
AT3-OVA Ctrl sgRNA
AT3-OVA Ptpn2 sgRNA
Tumor Size(mm²)
200
150
100
50
0
7 9 12 14 16 19 21 23 26 28 30 33
Days after tumor cell implantation

Fig. S12

***Figure S12. PTPN2 deletion does not repress the growth of AT3-OVA mammary tumors.*** AT3-OVA control cells (Ctl sgRNA) or those in which PTPN2 had been deleted by CRISPR RNP (*Ptpn2* sgRNA) were injected into the fourth inguinal mammary fat pads of female C57BL/6 mice and tumor growth was monitored.

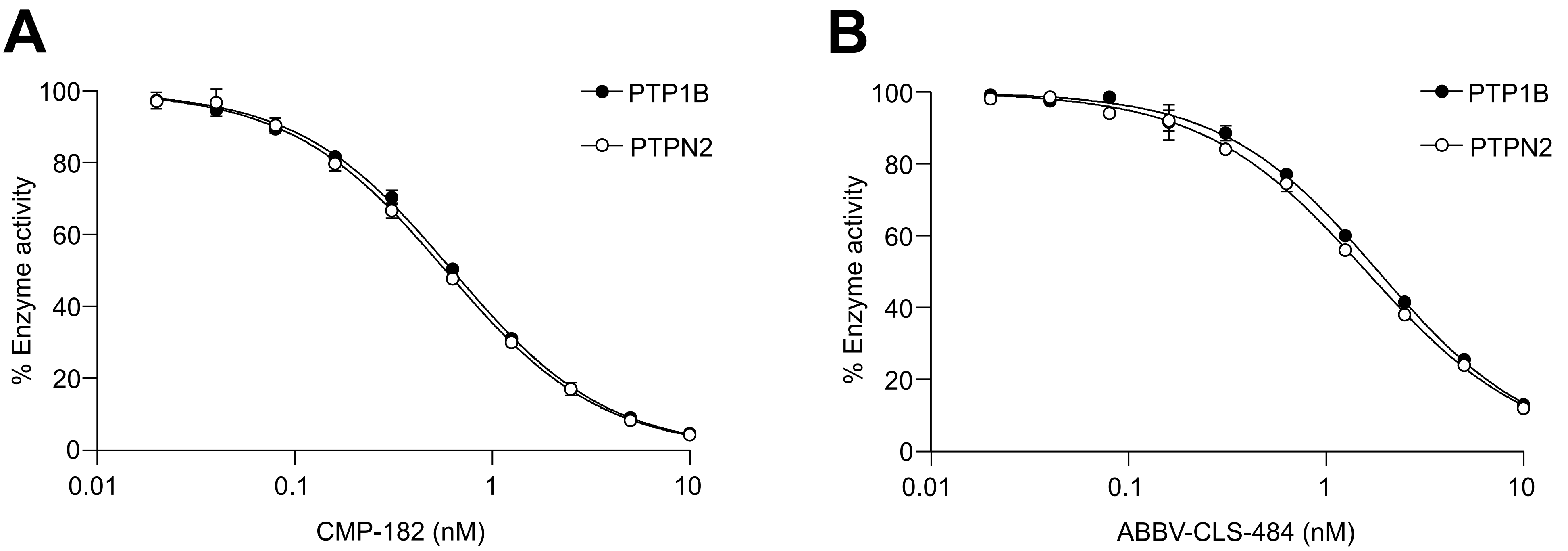
A
PTP1B
PTPN2
% Enzyme activity
100
80
60
40
20
0
0.01
0.1
1
10
CMP-182 (nM)
B
PTP1B
PTPN2
% Enzyme activity
100
80
60
40
20
0
0.01
0.1
1
10
ABBV-CLS-484 (nM)

**Fig. S13**

***Figure S13. Inhibition of PTP1B and PTPN2 by Compound 182 and ABBV-CLS-484.*** Effect of **a**) Compound 182 (CMP-182) versus **b**) ABBV-CLS-484 on PTP1B- and PTPN2-catalysed DiFMUP (10 μM) hydrolysis.
